# Titanium and platinum coatings reduce inflammation induced by gold on subretinal prosthesis

**DOI:** 10.64898/2026.07.13.737853

**Authors:** Mohajeet B Bhuckory, Vladimir Mamchik, Nicharee Monkongpitukkul, Davis Pham-Howard, Viktoryia Shautsova, Linh Mai Vu, Ludwig Galambos, Emma Butt, Keith Mathieson, Theodore Kamins, Daniel Palanker

## Abstract

Subretinal photovoltaic implants provide central vision to patients blinded by atrophic age-related macular degeneration, with acuity limited by their 100-µm pixels. Higher resolution requires smaller pixels incorporating three-dimensional electrodes, which can be fabricated by gold electroplating. However, the retinal response to exposed gold remains poorly characterized. Here, we evaluated gold biocompatibility on subretinal implants in Royal College of Surgeons rats and compared it with platinum- and titanium-coated surfaces. Although in-vivo optical coherence tomography revealed no overt structural disruption, gold implants induced cellular-scale anomalies, including abnormal morphology of rod bipolar cells, microglial accumulation near the implant, and increased cell death within days after implantation. These effects occurred across flat, pillar, and honeycomb geometries, indicating a material-rather than geometry-dependent response. By contrast, platinum- and titanium-coated implants showed substantially lower loss and morphological disruption of rod bipolar cells, together with markedly reduced microglial activation. These findings indicate that exposed gold surfaces can induce acute retinal inflammation and neuronal loss, whereas conformal platinum or titanium coatings substantially improve biocompatibility. Such coatings enable the development of three-dimensional subretinal prostheses with smaller pixels for improved visual resolution.

## Introduction

Loss of photoreceptors in diseases such as age-related macular degeneration (AMD) and retinitis pigmentosa (RP) leads to severe visual impairment, while the inner retinal circuitry remains largely preserved^1–3^. Subretinal prostheses offer a pathway to restoring vision by electrically stimulating these surviving neurons, primarily the bipolar cells in the inner nuclear layer (INL) and relying on transmission of their responses via retinal neural network to the brain. The photovoltaic subretinal implant PRIMA (Pixium Vision, Paris, France, now Science Corporation, California, USA) has demonstrated prosthetic central vision in patients with geographic atrophy secondary to AMD, with visual acuity of 20/420, which matches the sampling limit of its 100 μm pixel pitch^4–6^. Patients could also read smaller fonts using a digital zoom, albeit with a reduced field of view.

While these results represent a major advancement, higher visual acuity is required to expand the utility of retinal prostheses to a larger patient population. Pixels below 50 μm are required to exceed the legal blindness limit of 20/200. However, fundamental constraints on penetration of electric field limit the decrease of pixel size. This is particularly challenging with a degenerate human retina, where a subretinal debris layer frequently separates the implant from the INL by tens of micrometers^7^ (Fig.1 A). Such separation between electrodes and target neurons elevates the stimulation thresholds and limits the spatial resolution.

**Figure 1.**
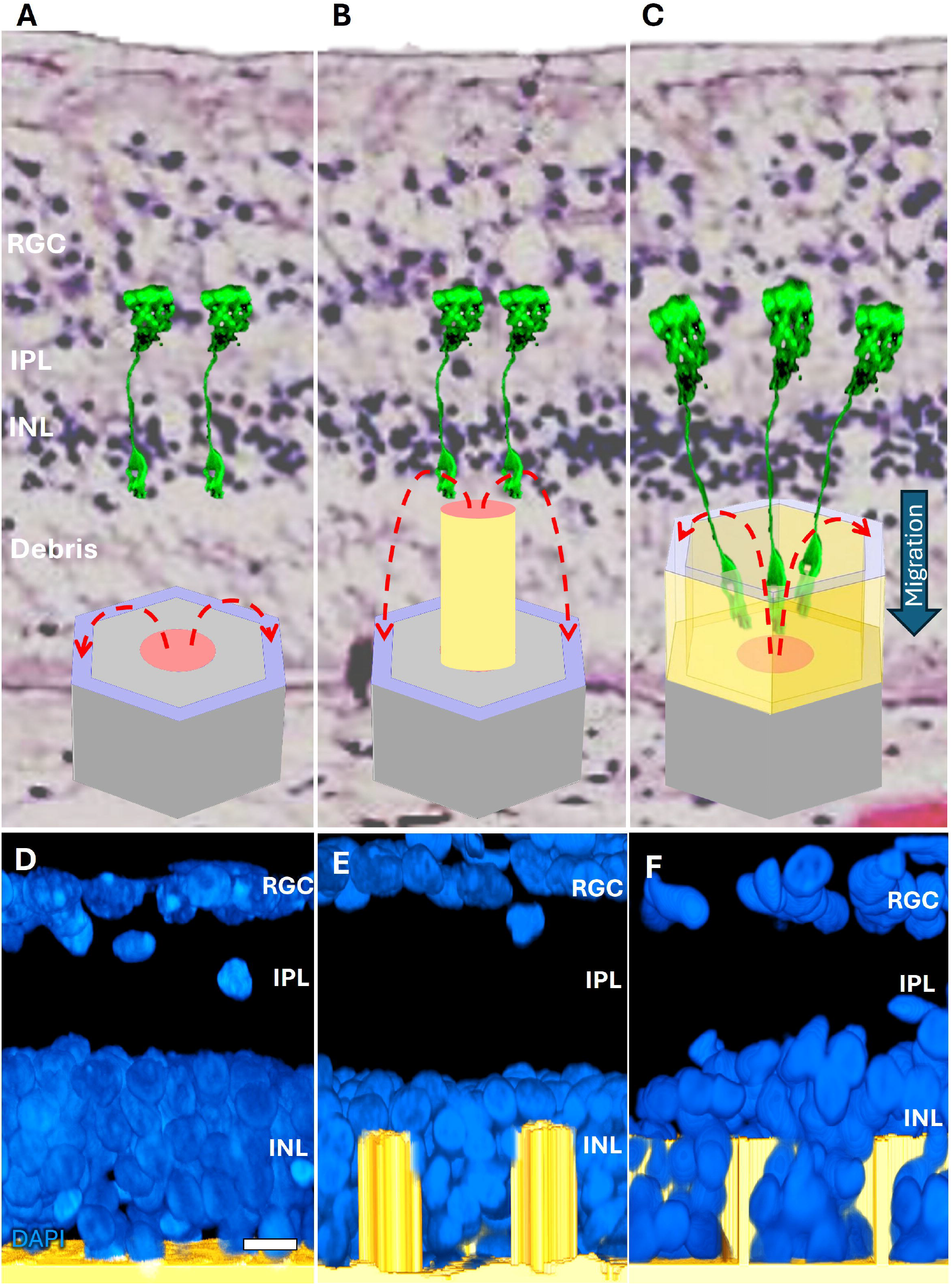
Three-dimensional subretinal prostheses improve proximity to target neurons within degenerated retina. (A) Schematic representation of a single bipolar photovoltaic pixel out of a planar array, similar to the PRIMA implant. The central active electrode (red) is surrounded by a return electrode (blue). In human degenerated AMD retina (overlaid histology), implant is separated from the inner nuclear layer (INL) by debris, limiting the penetration of electric field (red dash) to bipolar cells (green). (B) Pillars migrate through the debris layer and bring the active electrode closer to INL. (C) Within days, bipolar cells migrate into honeycombs, exposing them to strong electric field. (D-F) Confocal microscopy with DAPI staining of cell nuclei demonstrates integration of the three implant architectures with a degenerated retina in-vivo. (D) Flat implant. (E) Tissue migrating around the pillars (30 µm pitch) brings their tops into the middle of INL. (F) Migration of INL cells into the 30 µm wide honeycomb wells. Scale bar: 15 µm.

To overcome these limitations, three-dimensional (3D) electrode architectures have been developed to decouple pixel size from penetration depth of electric field (Fig. 1B-F). These include pillar electrodes penetrating up to INL (Fig. 1B) or honeycomb-shaped structures with elevated return electrodes that promote migration of retinal neurons into the wells, positioning them within the strong electric field (Fig. 1C). We have demonstrated successful integration of both architectures with a degenerated retina (Fig. 1 D-F) and efficient electrical stimulation^8–10^.

High-aspect-ratio conductive microstructures, such as pillars and honeycombs, can be electroplated in gold on top of photovoltaic arrays; a process compatible with a wafer-scale manufacturing^11^. Since specific capacitance of Au is substantially lower than that of SIROF, current injection occurs predominantly through the SIROF-coated electrode tops, which eliminates the need for electrical insulation of sidewalls, greatly simplifying the fabrication^11^. In contrast, electroplated platinum (Pt) exhibits stress-induced cracking, peeling, and deformation. Consequently, electroplated Au has become the structural foundation of the three-dimensional photovoltaic retinal prostheses.

Despite its widespread use in implants, ocular biocompatibility of Au remains poorly understood and contradictory in literature. Previous studies have reported highly variable results depending on the mode of administration, anatomical location, particle size, and duration of exposure^12–30^ (Table 1). Au-based nanoparticles have been associated with both therapeutic^18,22,27,31^ and inflammatory effects^17^, while implantation studies have reported outcomes ranging from good tissue tolerance to localized inflammatory responses^12–14^. Importantly, little is known about the reaction of bipolar cells to Au – neurons that are the most proximal to subretinal electrodes.

**Table 1.**
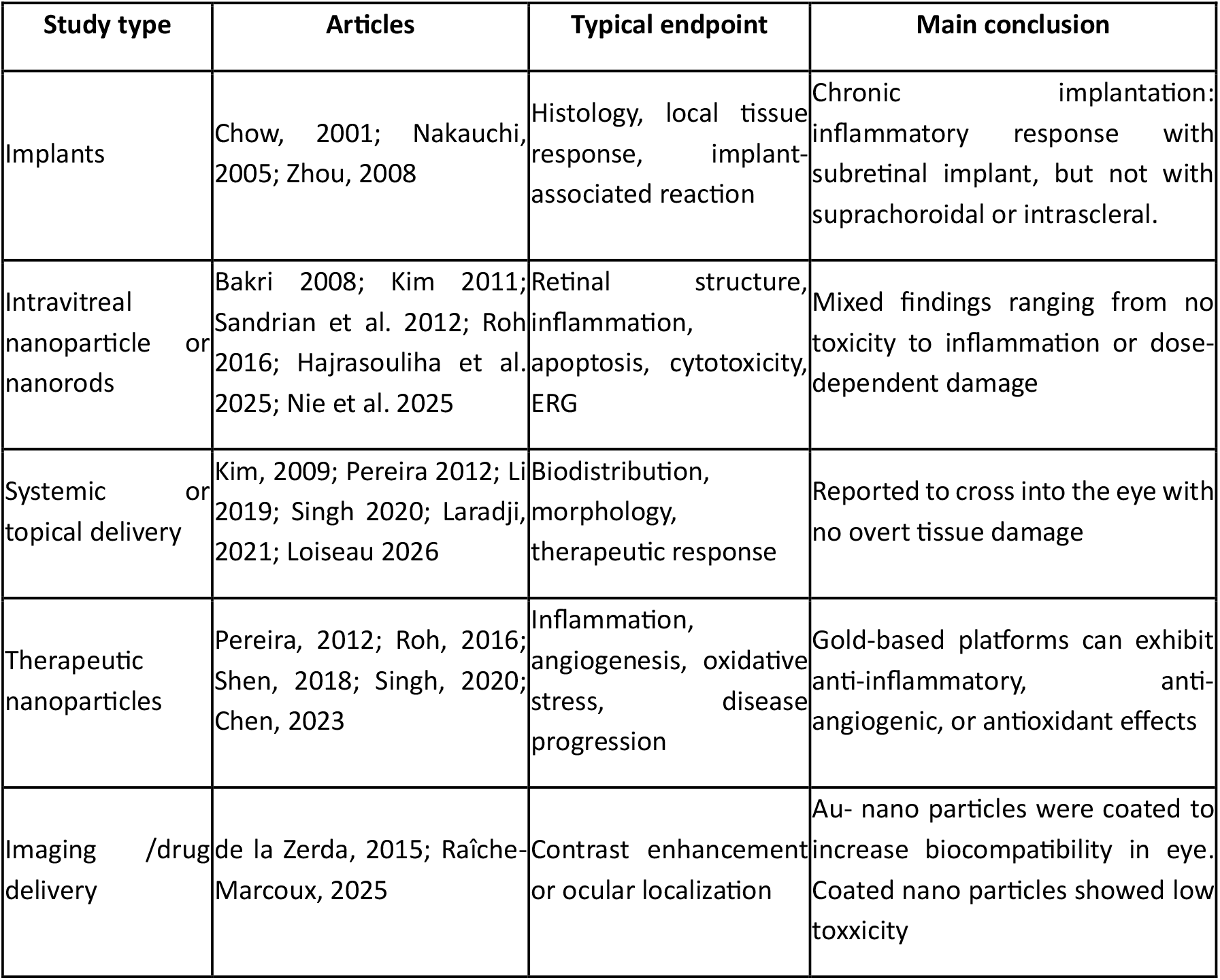
Literature review of gold biocompatibility or toxicity, with emphasis on the eye.

| Study type | Articles | Typical endpoint | Main conclusion |
| --- | --- | --- | --- |
| Implants | Chow, 2001; Nakauchi, 2005; Zhou, 2008 | Histology, local tissue response, implant-associated reaction | Chronic implantation: inflammatory response with subretinal implant, but not with suprachoroidal or intrascleral. |
| Intravitreal nanoparticle or nanorods | Bakri 2008; Kim 2011; Sandrian et al. 2012; Roh, 2016; Hajrasouliha et al. 2025; Nie et al. 2025 | Retinal structure, inflammation, apoptosis, cytotoxicity, ERG | Mixed findings ranging from no toxicity to inflammation or dose-dependent damage |
| Systemic or topical delivery | Kim, 2009; Pereira 2012; Li 2019; Singh 2020; Laradji, 2021; Loiseau 2026 | Biodistribution, morphology, therapeutic response | Reported to cross into the eye with no overt tissue damage |
| Therapeutic nanoparticles | Pereira, 2012; Roh, 2016; Shen, 2018; Singh, 2020; Chen, 2023 | Inflammation, angiogenesis, oxidative stress, disease progression | Gold-based platforms can exhibit anti-inflammatory, anti-angiogenic, or antioxidant effects |
| Imaging /drug delivery | de la Zerda, 2015; Raïche-Marcoux, 2025 | Contrast enhancement or ocular localization | Au- nano particles were coated to increase biocompatibility in eye. Coated nano particles showed low toxicity |

The present study evaluates biocompatibility of Au structures in subretinal space of RCS rats. We demonstrate that despite the absence of overt structural abnormalities in optical coherence tomography (OCT) imaging, chronic exposure to Au induces morphological changes in rod bipolar cells, microglial activation and increased cell death. We also demonstrate that Pt and Ti coatings strongly reduce such adverse biological responses, while preserving the structural and electrical advantages of 3D electrodes.

## Results

### Gold-induced retinal toxicity

*In-vivo* OCT of the degenerated RCS rat retina (Fig. 2A-D) displayed the expected absence of the outer nuclear layer, while preserved inner nuclear layer (INL) and the rest of the inner retina (Fig. 2A). At 6-9 weeks after implantation, the INL rests on top of the flat implants (Fig. 2B), gold pillar electrodes penetrated into INL (Fig. 2C) and honeycomb implants demonstrated stable retinal integration without overt structural abnormalities in OCT (Fig. 2D).

**Figure 2.**
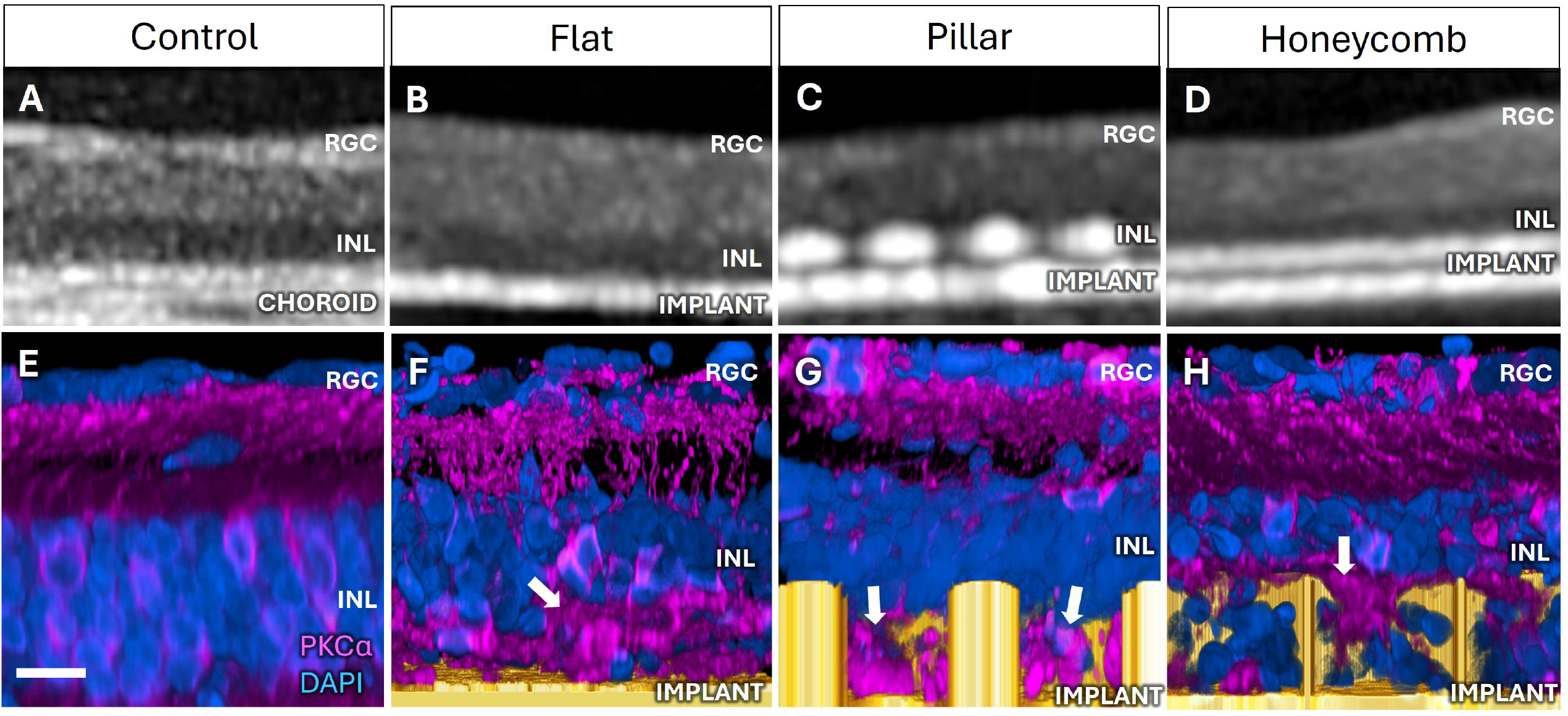
Subretinal prostheses six weeks post-implantation. (A–D) In-vivo OCT imaging of degenerated RCS retina. (A) Non-implanted control retina, demonstrating preservation of the inner retina after degeneration of photoreceptors. (B) Flat golden implants do not seem to affect the overlying retina. (C) Retina migrates between the 25µm-tall, 30 µm-pitch gold pillars, letting them penetrate into the INL. (D) Part of the INL migrates into the 40 µm-wide and 25 µm-deep gold honeycombs. OCT imaging revealed no overt structural abnormalities, indicating limited sensitivity to local cellular alterations associated with implants. (E-H) Confocal images with nuclei labeled with DAPI (blue) and PKCα-labeled rod bipolar cells (magenta) demonstrate presence of aberrant PKCα-positive clusters (white arrows) across all implant geometries. (E) Control area. (F) Flat golden implant. (G) Pillar electrodes. (H) Honeycomb structures. Scale bar: 20 µm.

However, on a microscopic level, rod BCs, labeled with PKCα in retinal wholemounts at six weeks post implantation, exhibited morphological abnormalities near the gold-coated implants of all three shapes: flat, pillar and honeycomb, compared to a non-implanted RCS retina (Fig. 2E-H). Aberrant clusters lacking associated nuclei were consistently observed adjacent to flat implants (Fig. 2F), pillars (Fig. 2G), and honeycombs (Fig. 2H). Reproducibility of this phenotype across various geometries suggests that it is associated with the implanted material.

To investigate the cellular response in greater detail, retinal wholemounts extracted together with integrated Au honeycombs were analyzed by confocal imaging of several immunofluorescent markers. PKCα labeling of rod bipolar cells revealed distinct abnormalities: in addition to normal membrane-associated PKCα staining surrounding the DAPI-positive nuclei (green arrows in Fig. 3A), implanted regions contained prominent PKCα-positive clusters lacking associated nuclei (white arrows). These aberrant accumulations differed morphologically from the normal bipolar cell architecture observed in non-implanted control retina (Fig. 3B). Surprisingly, quantification of the total PKCα-positive area showed no significant difference between honeycomb implants and controls (p= 0.69; Fig. 3C), indicating that amount of fluorescence alone did not reflect the profound morphological changes. In contrast to rod bipolar cells, secretagogin-positive cone bipolar cells maintained normal cellular morphology as in non-implanted controls (Fig. 3D-F, p= 0.12).

**Figure 3.**
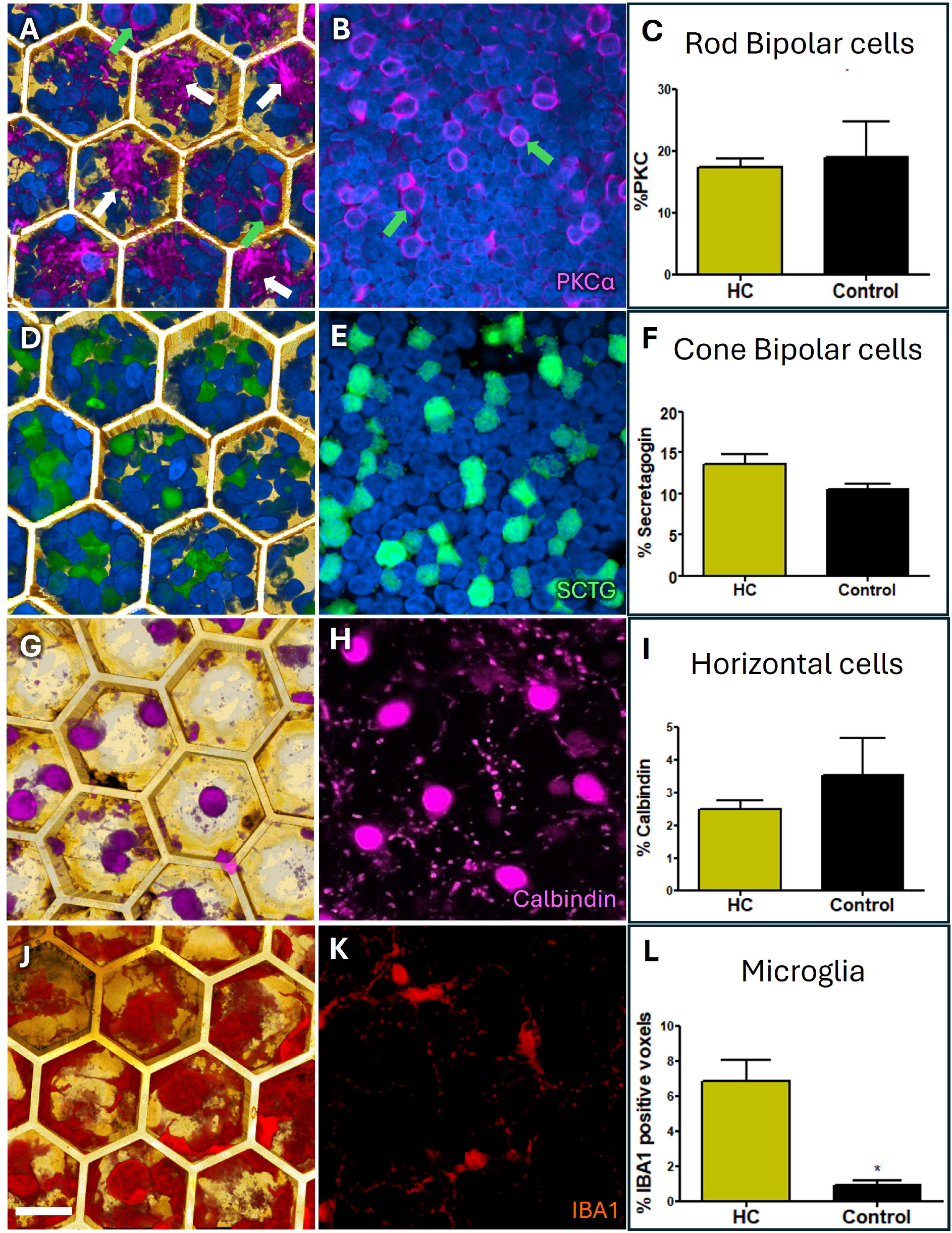
Immunohistochemical analysis of retinal cells above the implants. Confocal images of PKCα-labeled rod bipolar cells (magenta) in the retina with gold honeycomb implants (A) and a non-implanted control RCS retina (B). Normal membrane-associated PKCα staining surrounding DAPI-positive nuclei is indicated by green arrows, whereas aberrant PKCα-positive clusters lacking associated nuclei are indicated by white arrows. (C) No significant difference in PKCα-positive pixels between implanted and control groups despite prominent morphological abnormalities (ns, p = 0.69). Secretagogin (SCTG) immunolabeling of cone bipolar cells (green) in implanted (D) and control retina (E), demonstrating preserved morphology following implantation. (F) Number of Secretagogin-positive pixels is not significantly different between groups (ns, p= 0.12). Calbindin immunolabeling of horizontal cells (magenta) in implanted (G) and control retina (H). Horizontal cell numbers, morphology and processes remained preserved within honeycomb structures (I), with no significant difference between groups (ns, p= 0.10). IBA1 immunolabeling of microglia (red) in implanted retina (J) and control retina (K). Significantly increased microglial density in implanted retina compared to controls (*p=0.01, L). Scale bar: 20 µm.

Horizontal cells, another neuronal population in direct contact with the implant interface, were evaluated using calbindin immunostaining (Fig. 3G-I). They also exhibited normal cellular morphology and numbers within the honeycomb wells, comparable to those observed in a control retina (Fig. 3G-I p= 0.1). Together, these findings suggest that the retinal alterations induced by Au are selective to rod BCs rather than globally neurotoxic across all inner retinal cell populations.

IBA1 immunolabeling revealed marked accumulation of microglia within and around the wells of the honeycomb structures (Fig. 3J), substantially exceeding their sparse distribution in a control retina at the corresponding retinal depth (Fig. 3K-L, p= 0.01), indicating an inflammatory response associated with the implanted Au structures.

Collectively, these data demonstrate that although degenerated retina integrates with implanted golden 3D structures, abnormal morphology of rod bipolar cells and elevated microglial activation motivate investigation of biocompatible coatings.

### Superior biocompatibility of Pt and Ti

To investigate alternative coatings, we compared the retinal response to similar implants coated with platinum (Pt) or titanium (Ti). These materials were selected because they can be selectively electroplated onto side walls of 3D electrodes in a manner compatible with the rest of the implants manufacturing. With Au implants, retinas exhibited prominent PKCα-positive clusters localized at the implant-retina interface (Fig. 4A, white arrows), representing 22% of the total PKCα-positive pixels. In contrast, with Pt- and Ti-coated implants, clusters represented less than 5% of positive pixels (Fig. 4B-D). No significant difference was observed between Pt and Ti groups (p= 0.32, Figure 4D).

**Figure 4.**
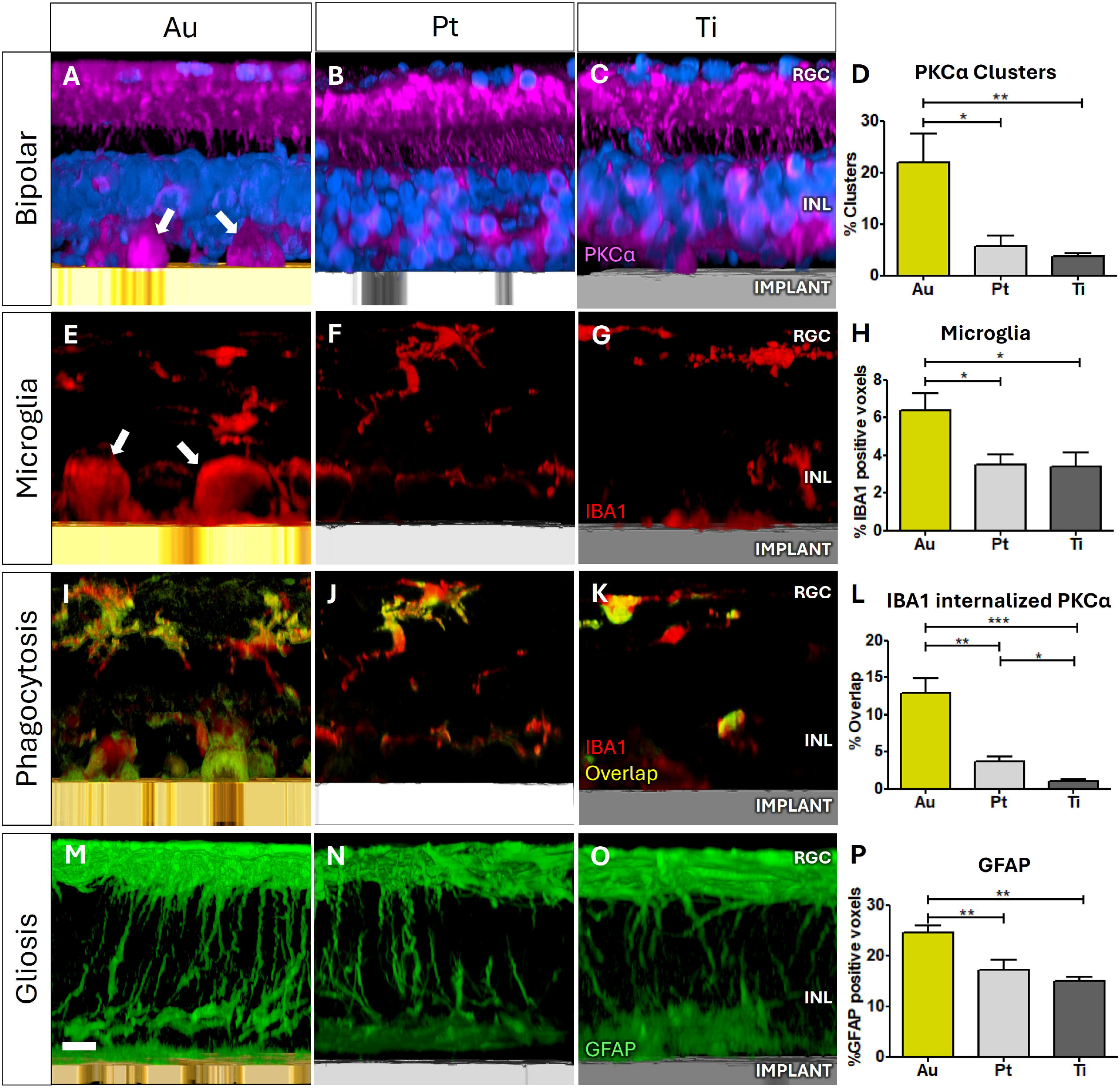
Platinum and titanium coatings reduce toxicity compared to gold implants. Representative confocal images of PKCα-labeled rod bipolar cells (magenta) in degenerated RCS retina with Au (A), Pt (B), or Ti(C) coated implants. White arrows indicate aberrant PKCα-positive clusters at the implant-retina interface. Retina with Pt and Ti implants has markedly fewer clusters than with Au. (D) Cluster area expressed as a percentage of total PKCα signal with Au, Pt and Ti implants. One-way Anova with Bonferroni post-hoc test: Au vs Pt: P = 0.01 (*), Au vs Ti: P = 0.005 (**), Pt vs Ti: P =0.32 (ns). (E-G) Representative confocal images of IBA1-positive microglia (red) adjacent to Au (E), Pt (F), and Ti (G) implants. Extensive microglial accumulation at the Au implant interface, with the formation of apparent giant immune cells, whereas much less inflammatory cell recruitment with Pt and Ti implants. (H) Quantification of microglia density: (*) Au vs Pt p= 0.01; (*) Au vs Ti p=0.01; (ns) Pt vs Ti p =0.89. (I-K) Co-localization between PKCα-positive bipolar cell material and IBA1-positive microglia in retinas implanted with Au (I), Pt (J) and Ti (K) structures. Overlapping signal indicates phagocytotic activity as internalization of bipolar cell-derived material by microglia. (L) PKCα signal internalized within IBA1-positive cells is much higher with Au implants compared to Pt and Ti. (**) Au vs Pt p= 0.006; (***) Au vs Ti p=0.0006; (*) Pt vs Ti p= 0.01. (M-O) GFAP immunolabeling in retinas implanted with Au (M), Pt (N), and Ti (O) structures. P) Significantly higher GFAP in the Au group compared to Pt and Ti at six weeks post-implantation. (**) Au vs Pt: p = 0.005; (**) Au vs Ti: p = 0.001; (ns) Pt vs Ti: p = 0.64. Scale bars: 20 µm.

Accumulation of microglial cells exhibiting inflammation marker IBA1 was observed adjacent to Au implants, frequently forming large aggregates resembling ‘giant cells’ (Fig. 4E). In contrast, Pt and Ti implants demonstrated reduced microglial recruitment (Fig. 4F, G), with significantly lower microglial density than with Au (Au vs Pt p= 0.01; Au vs Ti p=0.01), while no significant difference between Pt and Ti implants (Pt vs Ti p =0.89, Fig. 4H).

Co-localization (yellow) between PKCα-positive clusters and IBA1-positive microglia (red) is a measure of phagocytic internalization (Fig. 4I-L). Au implants demonstrated extensive overlap between PKCα and IBA1 labeling, indicative of bipolar cell-derived material contained within activated microglia (Fig. 4I). Much less overlap was observed in the Pt group (Fig. 4J), and even less than that with Ti implants (Fig. 4K,L: Au vs Pt p= 0.006; Au vs Ti p=0.0006, Pt vs. Ti p= 0.01).

Activation of microglia can promote Müller glial reactivity through the release of inflammatory mediators, leading to upregulation of the glial fibrillary acidic protein (GFAP), a well-established marker of retinal stress and injury^32^. Robust GFAP expression extending throughout the retinal thickness was observed with all metals (Fig. 4M-O), but more with Au than with Pt and Ti (Au vs Pt: p = 0.005; Au vs Ti: p = 0.001; Fig. 4P). Despite the general gliotic state of the degenerating retina, GFAP expression remains sensitive to the enhanced inflammatory response induced by gold six weeks post implantation.

Number of morphologically intact rod bipolar cells, identified by PKCα membrane staining surrounding a DAPI-positive nucleus, was reduced by approximately 24% with Au samples, relative to the non-implanted control retina, as opposed to 8% in Pt group and 4% with Ti implants (Fig. 5A-C): Au vs Pt p = 0.006; Au vs Ti p = 0.0002; Pt vs Ti p = 0.54 (Fig. 5D).

**Figure 5.**
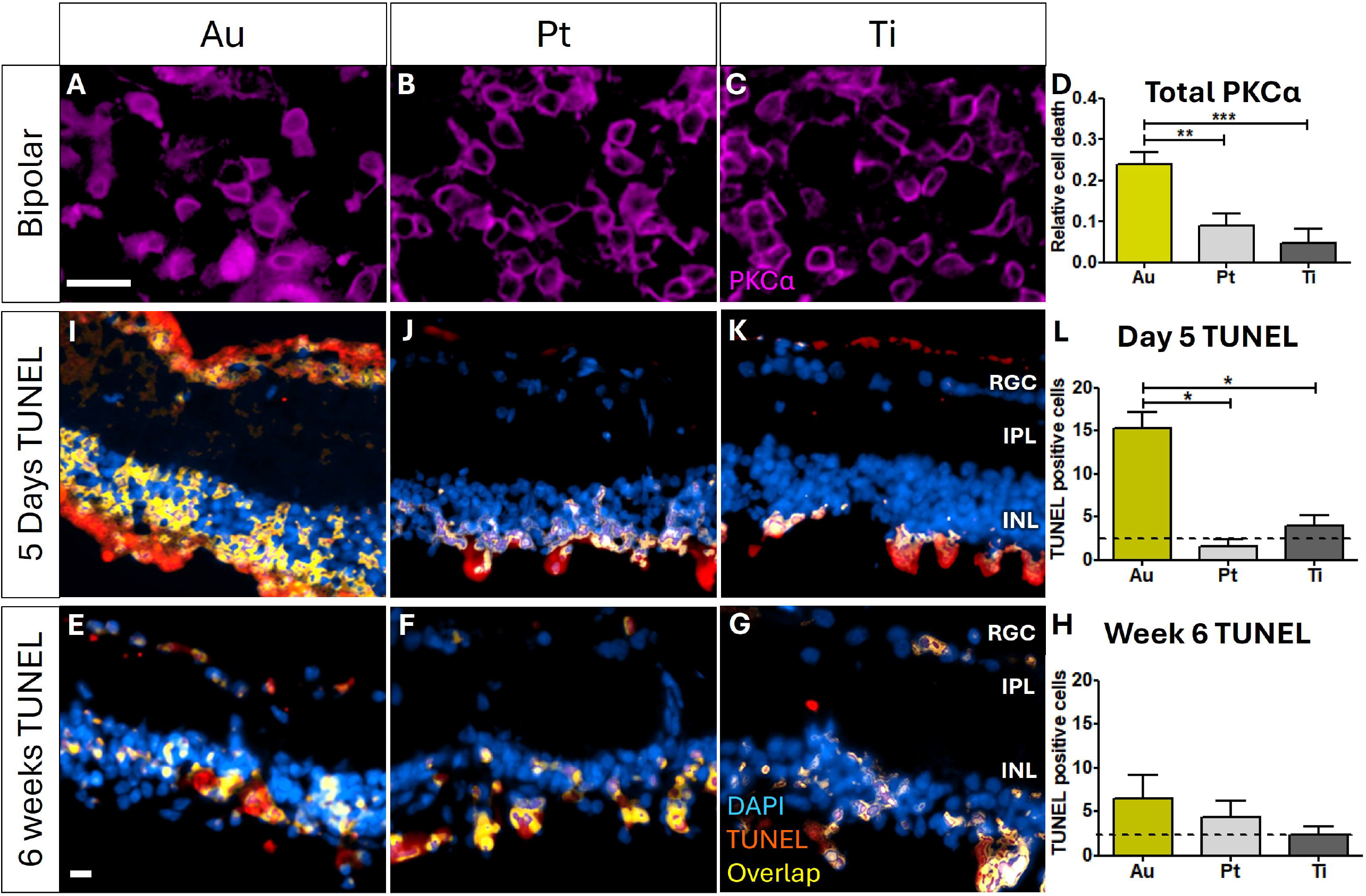
Apoptosis and loss of rod bipolar cells. (A-C) Top-down confocal images of PKCα-labeled rod bipolar cells (magenta) in degenerated RCS retina with Au (A), Pt (B), or Ti (C) implants. (D) Counting morphologically intact rod bipolar cells, defined as PKCα-positive membrane staining surrounding a DAPI-positive nucleus, shows significantly greater loss in the Au group compared with Pt and Ti. (**) Au vs Pt p = 0.006; (***) Au vs Ti p = 0.0002; (ns) Pt vs Ti p = 0.54. (I-K) TUNEL staining of retinal frozen sections five days after implantation with Au (I), Pt (J), or Ti (K) surfaces. (L) Overlap of TUNEL-positive nuclei and DAPI (labelled in yellow) demonstrates significantly increased cell death in the Au group compared with Pt and Ti groups, with no significant difference between Pt, Ti and the full control (dotted black line). (*) Au vs Pt p= 0.019; (*) Au vs Ti p= 0.042; (ns) Pt vs Ti p= 0.42. (E-G) TUNEL staining of retinal frozen sections six weeks after implantation with Au (E), Pt (F), or Ti (G). No significant differences between implanted groups or relative to non-implanted control levels, indicated by the dotted black line. (ns) Au vs Pt p = 0.54; (ns) Au vs Ti p = 0.16; (ns) Pt vs Ti p = 0.34. Scale bars: 20 µm.

### Timeline of cell death

To assess the timing of cell death, we compared apoptosis (TUNEL staining overlapping with DAPI-positive nuclei) at early (5 days) and late (6 weeks) time points. At 5 days post-op, Au samples exhibited increased apoptosis compared with Pt and Ti groups (Au vs Pt p= 0.019; Au vs Ti p= 0.042; Fig. 5I-K), with no significant difference between Pt and Ti (Pt vs Ti p= 0.42, Fig. 5L). However, at six weeks, no ongoing apoptotic cell death was detectable (Au vs Pt p = 0.54; Au vs Ti p = 0.16; Pt vs Ti p = 0.34; Fig. 5E-H); suggesting that an early wave of cell death ends by six weeks, resulting in a loss of rod bipolar cells and residual PKCα-positive debris or clustered structures at the implant interface.

Collectively, these findings demonstrate that Pt and Ti coatings significantly mitigate the loss and morphological abnormalities in rod bipolar cells, microglial activation, phagocytosis and macro-gliosis induced by Au implants.

## Discussion

Three-dimensional electrode architectures have emerged as a promising strategy for overcoming the pixel size limitations in retinal prostheses. Bringing stimulating electrodes closer to the inner retinal neurons using pillars or honeycomb structures can reduce the stimulation threshold, increase contrast, and thereby enable smaller pixels for higher visual acuity. Fabrication of these high aspect ratio structures utilizes gold electroplating, and gold is generally regarded as a biocompatible material widely used in medical implants.

Our study demonstrates that even with passive implants in subretinal space, Au induces loss and morphological abnormalities in rod bipolar cells and glial activation. Active Au electrodes may undergo dissolution under strong stimulation^12^ and produce toxic electrochemical products, such as hydrogen peroxide^33^. In advanced retinal degeneration, bipolar cells constitute a major component of the surviving inner retinal circuitry and represent the primary target of subretinal photovoltaic stimulation. Consequently, 24% loss of rod bipolar cells and other perturbations at the implant interface may have deleterious effects on prosthetic vision. We also demonstrate that Pt and Ti coatings isolating Au from the tissue exhibit much better biocompatibility.

Interestingly, only rod bipolar cells exhibited loss and abnormal morphology upon exposure to Au, whereas cone bipolar cells and horizontal cells remained largely unaffected. The precise origin of the PKCα-positive clusters remains to be determined, but persistence of these structures, together with the pronounced inflammatory response around Au implants, raises the possibility that cell death may involve mechanisms beyond classical apoptosis.

From a translational perspective, these findings have immediate implications for the development of the next-generation retinal prostheses. Au is uniquely suited for electroplating of high-aspect-ratio electrodes due to its conductivity, mechanical stability and compatibility with established microfabrication processes. Since Pt and Ti can be selectively electroplated on top of gold without affecting the rest of the implant, they offer a practical biocompatible solution, compatible with the wafer-scale implant manufacturing prostheses.

In conclusion, this study identified electroplated Au as a previously unrecognized source of toxicity to rod bipolar cells and offered a solution: Pt or Ti coating, which exhibits much better biocompatibility. More broadly, this study highlights the importance of evaluating interfaces between chronic implants and neural tissue, even when those materials are traditionally considered biocompatible and there are no macroscopic signs of toxicity.

## Materials and Methods

### Implant fabrication

Honeycomb implants used for assessment of biocompatibility were fabricated from crystalline silicon using deep reactive ion etching, as previously described^8,34^. Briefly, 40 µm hexagonal wells with 25 µm tall walls were patterned into 1mm diameter silicon substrates to create three-dimensional structures above a 30 µm thick silicon base. Implants were coated with a 200 nm layer of gold (Au) to prevent degradation of the underlying silicon in-vivo. For material comparison, similar sized flat implants were coated with 200 nm of Au or platinum (Pt) or titanium (Ti) by sputtering.

### Animals

All animal procedures were approved by the Stanford Administrative Panel on Laboratory Animal Care and adhered to the ARVO Statement for the Use of Animals in Ophthalmic and Vision Research. Experiments were performed in Royal College of Surgeons (RCS) rats obtained from a breeding colony maintained at Stanford University. Animals older than postnatal day 180 were selected to ensure a complete degeneration of photoreceptors prior to implantation.

### Subretinal implantation

Animals were anesthetized by intraperitoneal injection of ketamine (75 mg/kg) and xylazine (5 mg/kg). Following pupil dilation and preparation of the surgical field, a 1.5 mm scleral incision was created approximately 1 mm posterior to the limbus. A localized retinal detachment was generated by subretinal injection of sterile saline. Implants were inserted into the subretinal space and positioned at least 3 mm away from the retinotomy site. The conjunctiva was closed using 10-0 nylon sutures and topical antibiotic ointment was applied immediately after surgery. Retinal attachment and implant positioning and gross retinal morphology were verified by spectral-domain optical coherence tomography (OCT; Spectralis HRA+OCT, Heidelberg Engineering, Germany) before tissue collection.

For chronic studies, implants remained in the subretinal space for at least six weeks before tissue collection. For acute cell death studies, animals were euthanized five days after implantation. At the designated endpoint, animals were euthanized by intracardiac administration of pentobarbital/phenytoin solution. Eyes were immediately enucleated and rinsed in phosphate-buffered saline (PBS). Following removal of the cornea, iris, and lens, the posterior eyecup containing the implant was isolated. For whole-mount immunohistochemistry, a retinal region centered on the implant was carefully dissected and fixed in 4% paraformaldehyde at 4°C overnight.

### Immunohistochemistry and confocal imaging

Fixed retinal samples were permeabilized in PBS containing 1% Triton X-100 for 3 hours at room temperature and subsequently blocked with 10% bovine serum albumin (BSA). Samples were incubated with primary antibodies diluted in PBS containing 5% BSA and 0.5% Triton X-100 for 48 hours at room temperature. Following extensive washing in PBS containing 0.1% Tween-20, samples were incubated with fluorophore-conjugated secondary antibodies for 48 h at room temperature. Nuclei were counterstained with DAPI before final washes and mounting in antifade medium.

Primary antibodies included markers for rod bipolar cells (PKCα), cone bipolar cells (secretagogin), horizontal cells (calbindin), microglia (IBA1), and reactive Müller glia (GFAP). Antibody information is provided in Supplementary Table S1.

Retinal whole mounts were imaged using a Zeiss LSM 880 laser scanning confocal microscope (Carl Zeiss, Germany). Implant surfaces were visualized using reflected light. Z-stacks were acquired through the full retinal thickness extending from the inner limiting membrane to below the implant surface. Images were collected using a 40× oil immersion objective and processed using Fiji/ImageJ software. Identical acquisition settings were maintained for all experimental groups within each staining condition.

### Quantification

#### Clusters of rod bipolar cells

to quantify the PKCα-positive clusters, aberrant PKCα accumulations lacking associated DAPI-positive nuclei were manually outlined and measured using Fiji/ImageJ. Measurements were obtained from multiple fields per retina and averaged to generate a single value for each biological replicate.

#### Survival of rod bipolar cells

Intact cell density was determined by manually counting normal rod bipolar cells within a defined imaging area and normalizing them to the analyzed RCS control retinal surface area. Rod bipolar cells were identified as intact by the presence of PKCα-positive membrane staining surrounding a DAPI-positive nucleus. PKCα-positive clusters lacking nuclei were excluded from this count. Rod bipolar cell survival was expressed as the percentage of intact cells in the analyzed area relative to the number of intact cells in the equivalent control area.

#### Inflammatory and gliotic responses

Microglial activation was assessed by measuring IBA1-positive cell density within the retinal region directly adjacent to the implant. Reactive gliosis was quantified as a fraction of the area occupied by GFAP-positive staining within the analyzed field. Phagocytosis was assessed by overlap between PKCα-positive material and IBA1-positive microglia and quantified using colocalization analysis in Fiji/ImageJ. Internalized PKCα signal was expressed as a percentage of total PKCα cluster area overlapping with IBA1 staining.

#### Statistical analysis

Statistical analyses were performed using GraphPad Prism (version XX; GraphPad Software, San Diego, CA, USA). Data are presented as mean ± standard error of the mean (SEM). Comparisons among multiple groups were performed using one-way analysis of variance (ANOVA) followed by appropriate post hoc multiple-comparison testing. Differences were considered statistically significant when P < 0.05.

### TUNEL assay

Cell death was evaluated using terminal deoxynucleotidyl transferase-mediated dUTP nick-end labeling (TUNEL). Following euthanasia, implanted eyes were enucleated and fixed in 4% paraformaldehyde. Retinas were carefully separated from the implants, cryoprotected in graded sucrose solutions, embedded in optimal cutting temperature compound (OCT), and cryosectioned.

TUNEL labeling was performed according to the manufacturer’s protocol. Nuclei were counterstained with DAPI. Retinal cryosections were imaged with the confocal microscopy protocol as described earlier and processed using Fiji/ImageJ. Cells were considered TUNEL-positive only when TUNEL signal colocalized with a DAPI-positive nucleus. Tissue was collected after either 5 days or 6 weeks post implantation. TUNEL-positive nuclei were counted manually in retinal sections and normalized to the analyzed area in Fiji/ImageJ.

