## Supplemental Table 1 for "Titanium and platinum coatings reduce inflammation induced by gold on subretinal prosthesis"

Supplementary Table 1. Primary and secondary antibodies used for immunohistochemistry

| **Primary antibodies** | | |
|---|---|---|
| **Cell type** | **Host/Antibody** | **Supplier/Catalogue number** |
| Rod bipolar | Mouse/PKCα | SCBT/sc-8393 |
| Cone bipolar | Rabbit/Secretagogin | Cellsignaling/ 14037 |
| Horizontal cell | Mouse/Calbindin | Swant/ CB300 |
| Muller cell activation and astrocytes | Goat/Glial Fibrillary Acidic Protein (GFAP) | SCBT/SC-6170 |
| Microglia | Rabbit/IBA 1 | WAKO/ 019-19741 |
| **Secondary antibodies** | | |
| **Host/Reactivity** | **Fluorophore** | **Supplier/Catalogue number** |
| Donkey/Rabbit | AF488 | Thermofisher/ A-21206 |
| Donkey/Mouse | CY3 | Jackson lab/ 715-165-150 |
| Donkey/Goat | AF594 | Thermofisher/ A-11058 |
